# Active protein neddylation or ubiquitylation is dispensable for stress granule dynamics

**DOI:** 10.1101/418848

**Authors:** Sebastian Markmiller, Amit Fulzele, Reneé Higgins, Gene W. Yeo, Eric J Bennett

**Affiliations:** Department of Cellular and Molecular Medicine, University of California, San Diego, La Jolla, CA 92093, USA; Stem Cell Program, University of California, San Diego, La Jolla, CA 92093, USA; Institute for Genomic Medicine, University of California, San Diego, La Jolla, CA 92039, USA; Cell and Developmental Biology, Division of Biological Sciences, University of California, San Diego, La Jolla, CA 92093, USA

## Abstract

Many protein homeostasis stressors induce the formation of membraneless cytoplasmic stress granules (SGs) that contain large assemblies of repressed mRNAs and associated RNA binding proteins. Similar stressors have been shown to globally alter the function of the ubiquitin proteasome system (UPS) resulting in the accumulation of ubiquitylated proteins. Previous studies have demonstrated that ubiquitin and specific UPS components co-localize with SGs and that reducing the abundance or activity of ubiquitin pathway proteins can inhibit SG formation. These studies suggest that SG dynamics and composition may be regulated by ubiquitylation of SG resident proteins. Using ubiquitin-specific proteomic approaches, we demonstrate that many proteins, including some SG proteins are dynamically ubiquitylated upon SG-inducing sodium arsenite treatment. We utilized potent and selective inhibitors of the ubiquitin activating enzyme (UAE) or the NEDD8 activating enzyme (NAE) to directly test if active protein ubiquitylation or neddylation was required for SG dynamics. Using ubiquitin-site specific proteomics, we establish that UAE inhibition results in the rapid loss of nearly all protein ubiquitylation regardless of ubiquitin chain type. Addition of UAE or NAE inhibitors to cells did not alter arsenite-induced SG formation or dissolution. While we confirmed that ubiquitin co-localizes with both sodium arsenite and thapsigargin-induced SGs, antibodies that recognize all forms of ubiquitin more strongly co-localize with SGs compared to antibodies that preferentially recognize polyubiquitin or specific polyubiquitin-linkages. Interestingly, ubiquitin itself co-localizes with SGs in a UAE independent manner suggesting that the ubiquitin present within SGs is likely unconjugated ubiquitin. Our findings clearly demonstrate that active protein ubiquitylation or neddylation is not required for SG dynamics. These results suggest that ubiquitin-binding SG proteins may recruit free ubiquitin into SGs to modulate SG protein interactions.

## Introduction

Protein homeostasis can be disrupted by a diverse array of intrinsic and extrinsic stressors. Cellular insults such as oxidative and heat stress that globally disrupt protein folding result in both the accumulation of ubiquitylated proteins, and the induction of membraneless stress granules (SGs)^1,2^. SGs are enigmatic cellular structures that are comprised of translationally repressed mRNAs associated with a variety of RNA-binding proteins^3^. While the cellular function of SGs remain unclear, SG formation and SG resident proteins have been linked to human neurological disorders including amyotrophic lateral sclerosis (ALS) and frontotemporal degeneration (FTD)^3,4,5^. Further, pharmacological or genetic targeting of SG-resident proteins have been demonstrated to delay disease progression in models of ALS and Alzheimer’s disease^6-9^. These findings have fueled studies aimed at identifying SG RNA and protein constituents as potential modulators of SG dynamics and potential disease modifiers.

Genomic and proteomic characterization of both the SG RNA and protein constituents have revealed a marked compositional diversity in both stress granule proteins and RNAs^8,10,11^. Further, this diversity is altered by the exact stressor used to induce SGs as well as the cell type in which SGs are examined^8^. Examination of SG proteomes has revealed that proteins involved in regulating distinct post-translational modifications (PTMs) are often enriched within SGs. These findings suggest that PTMs may either regulate global SG dynamics or the recruitment of individual proteins into SGs^12^. For example, poly(ADP)-ribosylation was shown to regulate the SG localization and biophysics of the well-characterized SG protein TDP43, and ablation of the poly(ADP)-ribosylating enzyme, tankyrase, mitigated TDP43-assocaited toxicity *in vivo*^13^. These studies indicate that targeting PTMs may be an effective strategy to alter SG dynamics.

Numerous lines of evidence have implicated protein ubiquitylation or other ubiquitin-like modification systems, like neddylation, as a potential regulators of SG dynamics. First, components of the ubiquitin-proteasome system (UPS), including ubiquitin itself, have been shown to co-localize with SGs induced by a variety of protein homeostasis stressors^14-16^. Second, proteasome inhibition and the concomitant increase in polyubiquitylated proteins results in SG formation^15,17,18^. Third, genetic disruption or pharmacological inhibition of ubiquitin or neddylation components can disrupt SG dynamics in both *S. cerevisiae* and mammalian cells^14,16,18-23^. Despite this evidence, several key questions regarding the role of ubiquitylation in regulating SG dynamics remain unanswered. While ubiquitin has been shown to co-localize with SGs, whether polyubiquitylated proteins themselves, or proteins modified with specific ubiquitin-linkages are recruited to SGs is unknown. Further, SGs are variably induced by proteasome inhibition and sustained treatment with the proteasome inhibitor MG132 results in SG dissolution over time^15,17,18^. It is also unknown how many of the ubiquitin-system components that co-localize with SGs, require ubiquitin within SGs for their localization. The deubiquitylating enzyme USP10 is a well-characterized SG localized protein^21,24^. However, USP10 SG localization is determined by binding to another SG protein, G3BP1, and mutation of its active site, rendering it incapable of removing ubiquitin from substrates, had little impact on its localization or overall SG dynamics^22,25^. Despite the many links between the UPS and SGs, there has yet to be a demonstration that ubiquitylation of a specific SG protein is required for its SG localization, or that overall protein ubiquitylation or other ubiquitin-like protein modification pathways are needed to form or dissolve SGs.

Here, we directly examine the relationship between protein ubiquitylation and SG dynamics. Using sodium arsenite-induced oxidative stress to study SG dynamics, we show that arsenite induces polyubiquitylated protein accumulation and SGs with similar time scales and that arsenite-induced SGs co-localize with ubiquitin. Interrogation of global protein ubiquitylation using ubiquitin proteomics approaches revealed widespread alterations to the ubiquitin-modified proteome upon arenite-induced stress. Despite clear changes to some SG protein ubiquitylation, arsenite treatment did not result in global changes to known SG-resident protein ubiquitylation. Utilizing potent and specific inhibitors of either the ubiquitin activating enzyme (UAE) or the Nedd8 activating enzyme (NAE), we demonstrate that active protein ubiquitylation or neddylation is dispensable for arsenite-induced SG formation or dissolution. Further, we demonstrate that free, unconjugated ubiquitin localizes to SGs in a UAE-independent manner. Taken together, our results clearly demonstrate that active protein neddylation or ubiquitylation is not required for SG dynamics and that unconjugated ubiquitin may play a role in regulating SG protein localization or overall SG structure.

## Results

### Arsenite treatment induces stress granule formation and elevated protein ubiquitylation

We utilized the well-established stress granule (SG) inducer sodium arsenite to begin to characterize the relationship between protein ubiquitylation and stress granule dynamics. We first established the dynamics of SG formation by performing a time course of arsenite treatment on HeLa cells as well as an arsenite washout and recovery time course to characterize the timing of SG dissolution. We utilized the well-established SG marker G3BP1 to follow SG dynamics by immunofluorescence. Consistent with previous results, SGs could first be visualized 30 minutes after arsenite treatment which persisted up to two hours (**Fig. 1A**). Arsenite treatment for 1 hour followed by its removal resulted in a progressive loss in SGs and a near full recovery of diffuse G3BP1 staining was observed 3 hours after arsenite washout (**Fig. 1B**). SG formation coincided with a global increase in polyubiquitylated proteins and the induction of phosphorylated eIF2*α* indicating that alterations in protein degradation and overall protein homeostasis occur at the same time as SG formation (**Fig. 1C**). Protein ubiquitylation also initially decreased 2 hours after arsenite removal coincident with a partial reduction of phospho-eIF2*α* levels compared to arsenite-treated cells prior to washout (**Fig. 1D**). However, the abundance of polyubiquitylated proteins increased 3 hours after recovery indicating ubiquitin proteasome system function may be impacted beyond the time scales necessary for SG dissolution. To validate previous results demonstrating ubiquitin co-localization with stress granules, we treated three different commonly used cell types with sodium arsenite and visualized SGs with G3BP1 immunofluorescence. We were able to readily observe ubiquitin at arsenite-induced stress granules in all three cell lines using a ubiquitin antibody that recognizes both unconjugated and conjugated forms of ubiquitin. (**Fig. 1E**). Combined, these results suggest that the timing of alterations to protein ubiquitylation largely mirror SG dynamics and ubiquitin co-localizes to stress granules indicating a possible role for protein ubiquitylation during SG formation or dissolution.

**Figure 1.**
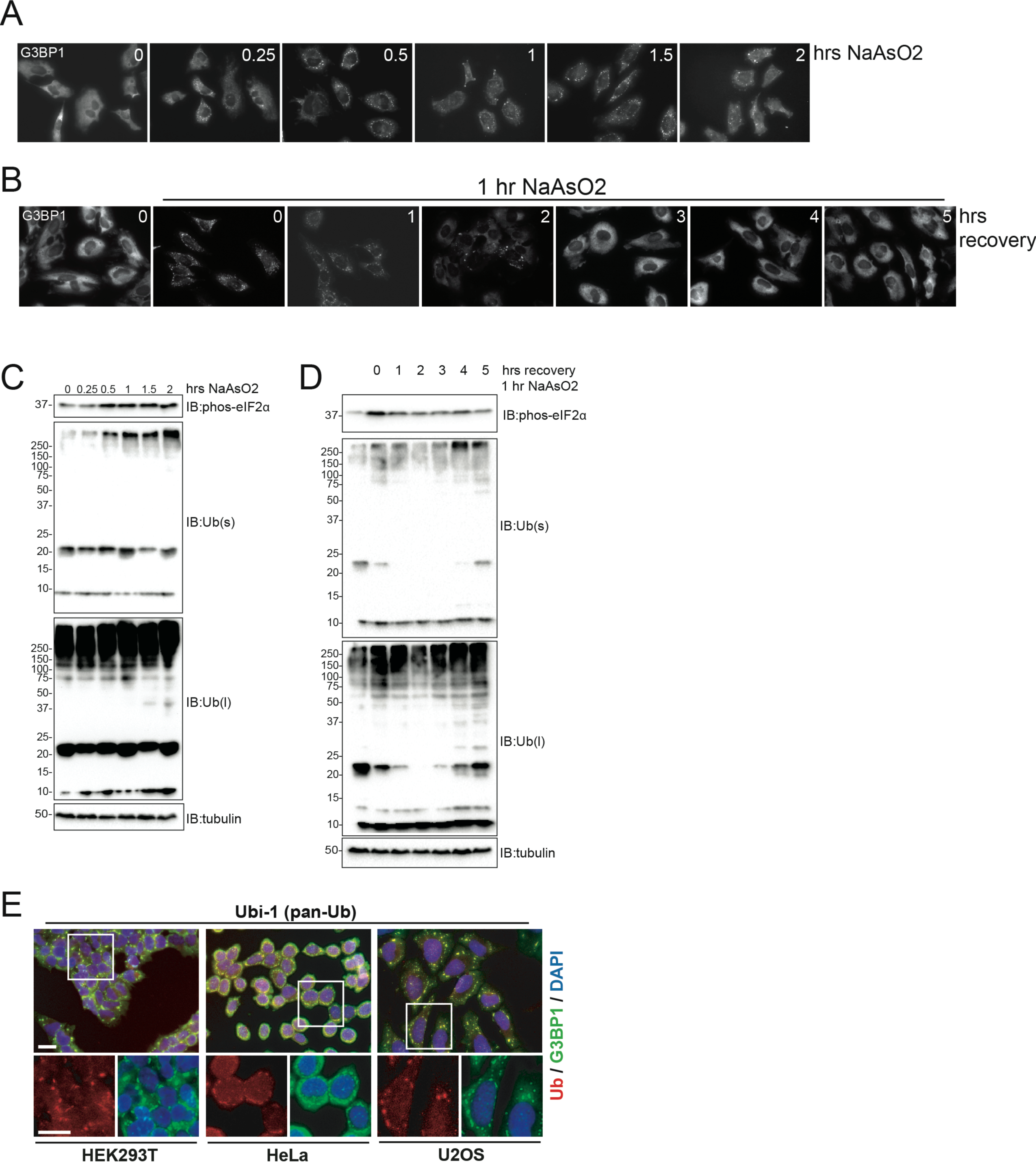
Sodium arsenite treatment induces stress granule formation and global protein ubiquitylation. A).HeLa cells were treated with sodium arsenite (500μM) over the indicated time course. Cells were fixed and stress granules were visualized using G3BP1 immunofluorescence at the indicated times. B).HeLa cells were either untreated or treated with sodium arsenite (500μM) for 1 hour. The sodium arsenite-containing media was replaced with fresh media without sodium arsenite and cells were allowed to recover for the indicated times. Stress granules were visualized using G3BP1 immunofluorescence. C,D) Hela whole cell lysates corresponding to the sodium arsenite treatment (C) and recovery (D) were separated by SDS-PAGE and immunoblotted as indicated. s and l denote short and long exposures, respectively. E) The indicated cell lines were treated with sodium arsenite (500μM for 1h). Cells were then fixed, and stress granules were localized by G3BP1 immunofluorescence or using G3BP1-GFP fluorescence for 293 cells (green). Ubi-1 antibody staining was used to localize ubiquitin (red). Scale bar = 20μm.

### Identification of resident SG protein ubiquitylation upon arsenite treatment

Our data, as well as previous reports, suggest that protein ubiquitylation may regulate overall SG dynamics and the ability of specific proteins to localize to SGs upon protein homeostasis stress. We utilized quantitative ubiquitin site-specific proteomic approaches to identify proteins whose ubiquitylation status was altered upon arsenite treatment^26, 27^. HeLa cells grown in media containing ^13^C^15^N-labeled lysine were either untreated or treated with arsenite for 20 or 45 minutes or were treated with arsenite for 45 minutes followed by washout and recovery for 3 hours. Consistent with our previous results, arsenite treatment resulted in a global increase in protein ubiquitylation which was reduced upon arsenite removal and recovery for 3 hours (**Fig. 2A,B, Table S1**). Arsenite treatment did not result in overt changes to protein abundance, at least for the depth of proteome coverage achieved in this experiment. Greater than 35% of all ubiquitin-modified peptides quantified from cells treated with arsenite for 45 minutes increased or decreased in abundance more than 2-fold indicating that a significant fraction of protein ubiquitylation is impacted by arsenite treatment (**Fig. 2C**). Modest, but consistent abundance increases in all but lysine11(K11)-linked ubiquitin-chains were observed upon 45-minute arsenite treatment which were reduced during arsenite washout and recovery (**Fig. 2D**). Interestingly, K63-linked ubiquitin chains also increased upon arsenite treatment which suggests that regulatory protein ubiquitylation that does not target proteins for degradation may be involved in the response to arsenite-induced oxidative stress, and by extension, SG dynamics. We utilized a curated list of validated SG resident proteins to directly examine if known SG proteins were selectively ubiquitylated upon arsenite treatment (**Table S2**). The abundance of 102 ubiquitylated peptides across 43 known SG proteins was not significantly altered upon arsenite treatment or washout compared to untreated samples indicating that, at least taken together, SG resident protein ubiquitylation was not specifically altered during conditions that induce SG formation (**Fig. 2E**). However, examination of individual SG proteins revealed known SG proteins with multiple ubiquitylation events that either robustly increased (HNRNPK) or decreased (RPS3, PABPC4) upon arsenite treatment (**Fig. 2F**). Our data suggest that while SG protein ubiquitylation is not globally impacted by arsenite treatment, it remains possible that the ubiquitylation of specific SG proteins govern their SG localization.

**Figure 2.**
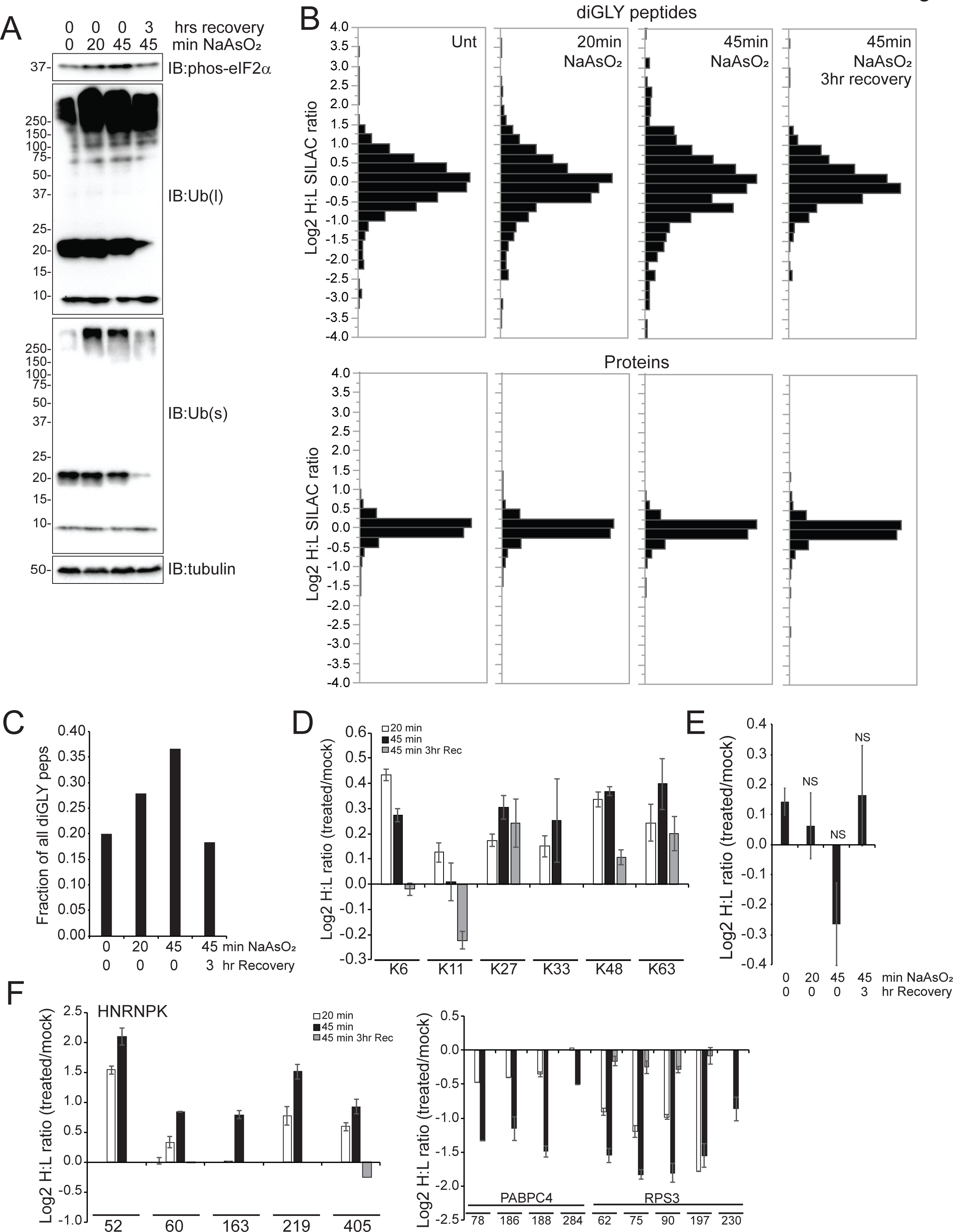
Stress granule protein ubiquitylation is largely unaffected by sodium arsenite treatment. A).Hela cells were treated with sodium arsenite (500μM) for the indicated times with or without washout and recovery. Whole cell extracts were separate by SDS-PAGE and immunoblotted as indicated. s and l denote short and long exposures, respectively. B).Heavy (^13^C^15^N) lysine-labeled Hela cells were treated with sodium arsenite (500μM) for the indicated times with or without washout and recovery. Corresponding untreated light-labeled Hela cells were mixed with heavy arsenite treated cells and whole cell lysates were generated and digested with trypsin. Ubiquitin-modified peptides (diGLY peptides) were immuno-isolated and both diGLY-modified peptides and total peptides were analyzed by mass spectrometry. The log2 heavy to light ratios are depicted for all diGLY-modified peptides (top) and total proteins (bottom) for each arsenite treated sample. C).The fraction of all quantified diGLY peptides that increased or decreased in abundance greater than 2-fold upon sodium arsenite treatment or washout. D).The log2 heavy to light ratio corresponding to diGLY-modified peptides from ubiquitin. The individual modified lysine that was quantified is indicated. Error bars denote SEM of multiple quantification events for a given peptide. E).The median log2 ratio of all diGLY-modified peptides quantified from known stress granule proteins is depicted for each sodium arsenite treated sample. Error bars denote SEM for all diGLY quantification events observed on known stress granule proteins. NS=not significant F).The log2 heavy to light ratio corresponding to diGLY-modified peptides from the stress granule proteins HNRNPK (left) or PABPC4 and RPS3 (right). The individual diGLY-modified lysine that was quantified is indicated. Error bars denote SEM of multiple quantification events for a given peptide.

### Acute pharmacological inhibition of the ubiquitin activating enzyme results in a rapid loss of protein ubiquitylation

Ubiquitin has been observed to co-localize with SGs and ubiquitin system components have been demonstrated to both localize to SGs and regulate SG dynamics implicating a role for protein ubiquitylation during SG formation or dissolution. However, a direct evaluation of whether active protein ubiquitylation is required to either form or resolve SGs has not been reported. To perform such an examination, we utilized a newly developed specific and potent ubiquitin E1 activating enzyme (UAE) inhibitor (TAK-243, also known as MLN7243) to acutely inhibit protein ubiquitylation^28^. We first determined the concentration and time dependence on the reduction of protein ubiquitylation upon TAK-243 (which we refer to as Ub-E1i) treatment. Addition of increasing amounts of Ub-E1i to HCT116 cells for 4 hours resulted in a dose-dependent decrease in polyubiquitylated proteins with complete abrogation of observable polyubiquitylated material with Ub-E1i treatment above 0.5μM (**Fig. 3A**). Proteasome inhibition resulted in the well-characterized increase in total protein ubiquitylation which was completely blocked upon co-treatment with Ub-E1i. Ub-E1i addition resulted in time-dependent decrease in polyubiquitylated material, and consistent with previous reports, Ub-E1i treatment was selective for UAE as cullin neddylation was unaffected after 4 hours of Ub-E1i treatment but was completely blocked by addition of the Nedd8 activating enzyme (NAE) inhibitor MLN4924 (TAK-924 which we refer to as N8-E1i) (**Fig. 3A,B**)^29^. Having established a concentration and time in which Ub-E1i treatment completely blocked protein ubiquitylation, as determined by immunoblotting, we set out to establish the impact of Ub-E1i inhibition on individual protein ubiquitylation using quantitative ubiquitin-site specific proteomics. Heavy (^13^C^15^N) lysine-labeled HCT116 cells were treated with Ub-E1i alone or in combination with MG132 for 4 hours and were mixed with unlabeled HCT116 cells that were either untreated or MG132 treated. MG132 treatment allows for a deeper characterization of the impact of Ub-E1i addition as proteasome inhibition significantly increases the abundance of ubiquitylated proteins. Total mixed lysates were digested with trypsin and diGly-modified peptides were enriched by immuno-affinity approaches. As expected, Ub-E1i treatment resulted in a robust reduction in the abundance of the clear majority of ubiquitylated peptides identified (**Table S3**). More than 80% of all quantified ubiquitin-modified peptides were reduced in abundance more than 1.7-fold with or without MG132 treatment (**Fig. 3C,D**). As observed with proteasome inhibition alone, UAE inactivation had little impact on overall protein abundance after 4 hours of treatment (**Fig. 3C, Table S3**). However, a more complete characterization of low abundance proteins would likely reveal robust protein abundance alterations. As would be predicted upon UAE inhibition, all detected ubiquitin-linkage peptides were reduced more than 8-fold, indicating that Ub-E1i treatment severely reduces total protein polyubiquitylation regardless of chain type (**Fig. 3E**). A direct examination of the ubiquitylation status of known SG proteins revealed that Ub-E1i treatment, like observed for more than 80% of all ubiquitylated peptides, resulted in a clear reduction of SG protein ubiquitylation across a diversity of individual ubiquitylation sites (**Fig. 3F**). Interestingly, a subset of diGly-modified peptides were refractory to UAE inactivation (**Fig. 3C,D**). These peptides either arise from neddylation or their ubiquitylation is not rapidly reversible upon UAE inactivation. Regardless, our results demonstrate that Ub-E1i treatment results in the rapid loss of greater than 80% of all ubiquitylation events.

**Figure 3.**
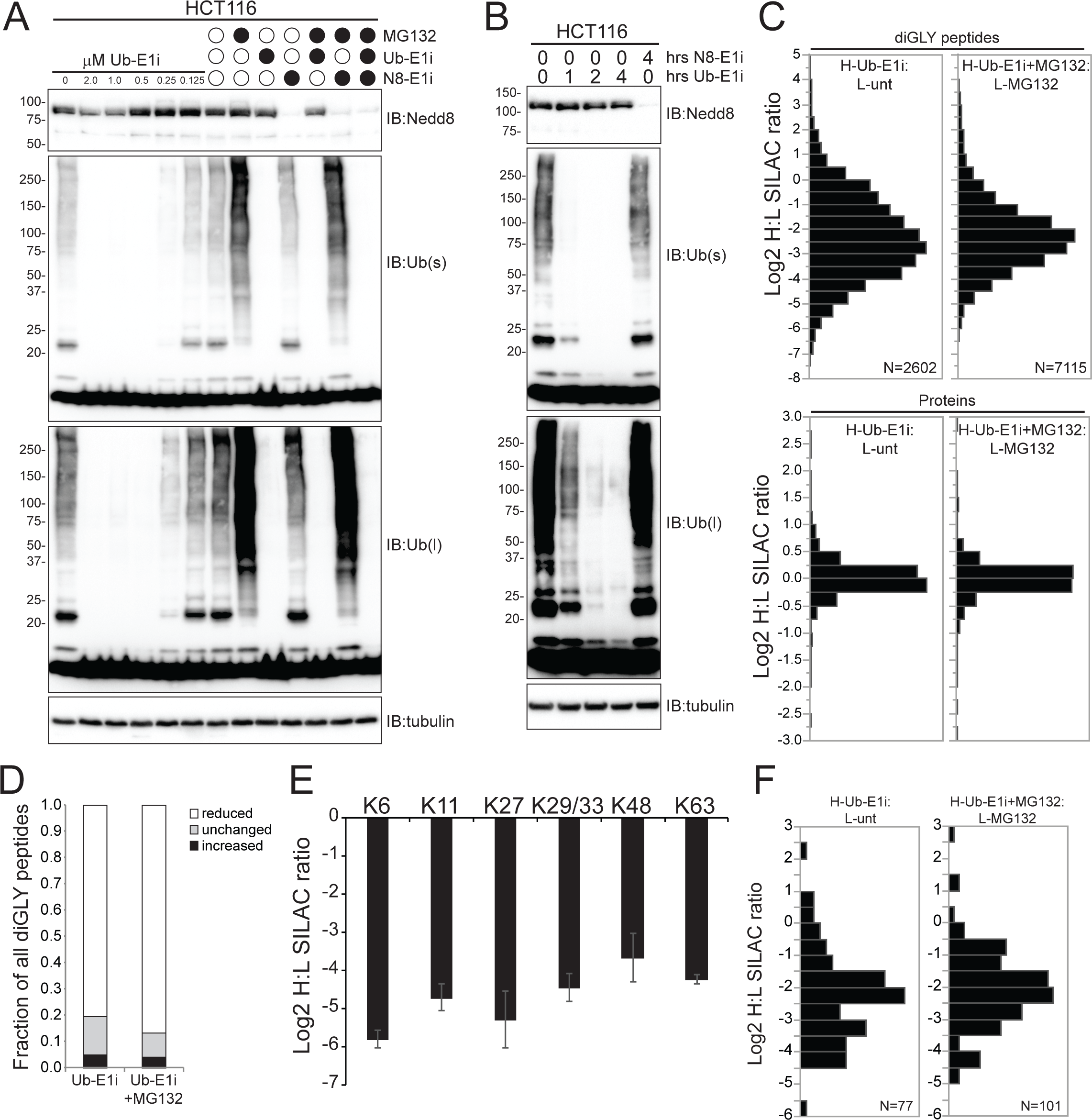
The ubiquitin activating enzyme inhibitor TAK-243 rapidly ablates ubiquitin conjugates in cells. A,B) HCT116 cells were treated with MG132, the ubiquitin activating enzyme inhibitor TAK-243 (Ub-E1i), or the NEDD8 activating enzyme inhibitor TAK-924 (N8-E1i) as indicated. Whole cell lysates were separated by SDS-PAGE and immunoblotted as indicated. s and l denote short and long exposures, respectively. C).Heavy (^13^C^15^N) lysine-labeled HCT116 cells were treated with Ub-E1i with or without MG132 for 4 hours. Corresponding untreated or MG132-treated light-labeled HCT116 cells were mixed with heavy-labeled Ub-E1i-treated cells and whole cell lysates were generated and digested with trypsin. Ubiquitin-modified peptides (diGLY peptides) were immuno-isolated and both diGLY-modified peptides and total peptides were analyzed by mass spectrometry. The log2 heavy to light ratios are depicted for all diGLY-modified peptides (top) and total proteins (bottom) for each sample. D).The fraction of all quantified diGLY-modified peptides that increased or decreased in abundance greater than 1.7-fold or were unaltered upon Ub-E1i treatment with or without proteasome inhibition. E).The log2 heavy to light ratio corresponding to diGLY-modified peptides from ubiquitin after Ub-E1i treatment alone. The individual ubiquitin-modified lysine that was quantified is indicated. Error bars denote SEM of multiple quantification events for a given peptide. F).The log2 ratio of all diGLY-modified peptides quantified from known stress granule proteins is depicted after Ub-E1i treatment with or without proteasome inhibition.

### Active protein ubiquitylation or neddylation is not required for SG formation or dissolution

Having established UAE inactivation via Ub-E1i treatment as a powerful tool to rapidly and robustly ablate protein ubiquitylation and probe the dependence of any pathway on active protein ubiquitylation, we set out to test if SG formation or dissolution required protein ubiquitylation. We first utilized a previously characterized 293T cell line expressing the well-established SG protein, G3BP1, tagged with GFP at its endogenous C-terminus using CRISPR/Cas9-based approaches^8^. Consistent with previous results, arsenite addition resulted in a rapid increase in SG formation as determined by G3BP1-GFP coalescence (**Fig. 4A,B**). Pre-treatment with Ub-E1i followed by treatment with two different concentrations of arsenite did not delay the kinetics of SG formation over a 2-hour time course (**Fig. 4A,B, S1A,B**). At 100μM arsenite, Ub-E1i addition resulted in a small enhancement of SG formation at early time points, indicating that the added stress of UAE inhibition may accelerate SG formation for lower concentrations of arsenite. Pre-treatment with the NAE inhibitor MLN4924 (N8-E1i) followed by arsenite treatment did not affect SG formation at either of the tested arsenite concentrations (**Fig. 4A,B, S1A,B**). Addition of either Ub-E1i or N8-E1i alone did not induce SG formation (**Fig. S1A**). Taken together, these observations indicate that neither active protein ubiquitylation nor neddylation are required for initial arsenite-induced SG formation. We then determined if protein ubiquitylation or neddylation was required for SG dissolution following arsenite removal. 293T G3BP1-GFP cells were pre-treated with DMSO, Ub-E1i, or N8-E1i for 1 hour and SGs were formed by treatment with arsenite in the presence of inhibitors for 1 hour. SG dissolution was monitored after arsenite washout in media that lacked Ub-E1i or N8-E1i. While NAE inhibition has no measurable impact on SG dissolution, Ub-E1i addition resulted in a minor delay in SG dissolution (**Fig. 4C,D**). We validated these results in HeLa cells using G3BP1 antibodies to monitor SG formation and clearance. As observed in 293T cells, UAE inhibition did not delay the kinetics of arsenite-induced SG formation using two different arsenite concentrations, with a slight acceleration of SG formation at early time points (**Fig. 4E,F, S1C**). We were unable to validate the small delay in SG dissolution upon Ub-E1i treatment seen in 293T cells as UAE inactivation had no impact on SG clearance in HeLa cells (**Fig. 4G,H, S1D**). Taken together, our results clearly indicate that acute inhibition of protein ubiquitylation or neddylation has little to no impact on SG dynamics.

**Figure 4.**
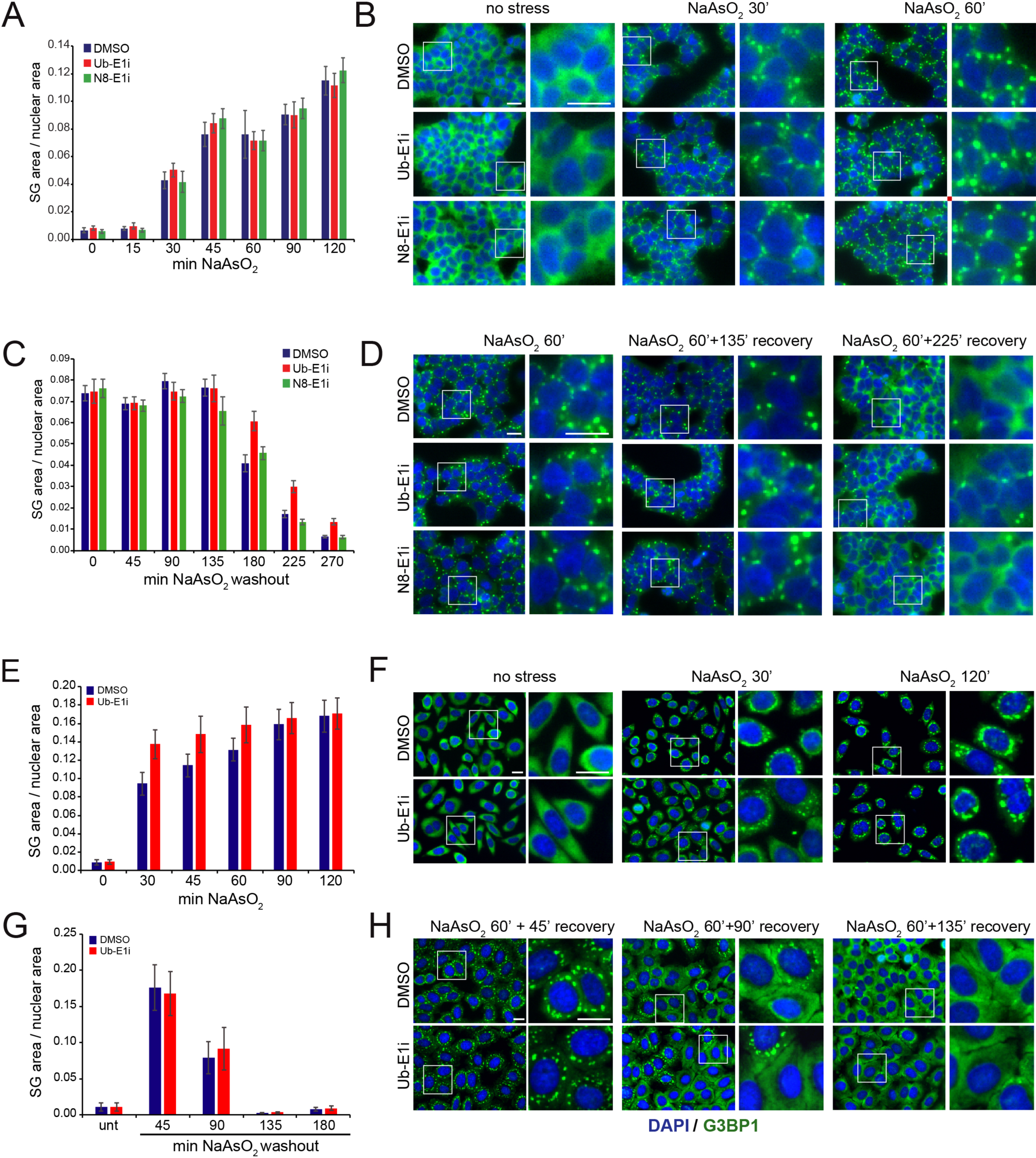
Stress granule dynamics are unaffected by inhibition of protein ubiquitylation or neddylation. A).293T cells expressing G3BP1-GFP were pre-treated 60’ with DMSO (blue bars), Ub-E1i (1μM, red bars), or N8-E1i (1μM, green bars) followed by treatment with sodium arsenite (500μM) for the indicated times. Cells were fixed and imaged at the indicated times. The stress granule and nuclear area was quantified using custom image analysis scripts for the indicated time points and treatments. Error bars depict standard deviation of the mean. B).Representative images of G3BP1-GFP 293T cells treated with or without sodium arsenite and with or without Ub-E1i or N8-E1i pre-treatment. C).G3BP1-GFP 293T cells were pre-treated with DMSO, Ub-E1i, or N8-E1i for 60’ and subsequently treated with or without sodium arsenite (500μM) for 60’ at which time the sodium arsenite and UAE or NAE inhibitors were washed out and cells were fixed and imaged at the indicated times following sodium arsenite washout. The stress granule and nuclear area was quantified using custom image analysis scripts for each condition. D).Representative images for the indicated time points and treatments quantified in panel C. E).Hela cells were pre-treated with DMSO or Ub-E1i for 90’ followed by treatment with sodium arsenite (500μM) for the indicated times. Cells were fixed, SGs were localized using G3BP1 immunofluorescence, cells were imaged, and SG and nuclear area was quantified at the indicated times. F).Representative images for the indicated time points and treatments quantified in panel E. G).Hela cells were pre-treated with DMSO or Ub-E1i for 90’ followed by treatment with sodium arsenite (250μM) for 60’ at which time the sodium arsenite and Ub-E1i was washed out and cells were fixed and imaged at the indicated times following sodium arsenite washout. Cells were fixed, SGs were localized using G3BP1 immunofluorescence, cells were imaged, and SG and nuclear area was quantified at the indicated times. Error bars depict standard deviation of the mean. H).Representative images for the indicated time points and treatments quantified in panel G. Error bars in panels A,C,E and G depict standard deviation of the mean. Scale bars in panels B,D,F and H = 20μm in all panels.

### Unconjugated ubiquitin co-localizes with SGs in a UAE independent manner

In accordance with previous studies, we were able to show that ubiquitin co-localizes with SGs using immunofluorescence techniques (**Fig. 1E)**. Because ubiquitin exists in a variety of biochemically distinct pools within cells (e.g. unconjugated “free” ubiquitin and lysine-48 linked polyubiquitin chains)^30^, we set out to carefully characterize which forms of ubiquitin specifically co-localize with SGs. As detailed earlier, sodium arsenite treatment in three cell types resulted in robust co-localization of SGs with ubiquitin using a pan-ubiquitin antibody that recognizes both unconjugated and conjugated ubiquitin (**Fig. 1A**). Staining with two different ubiquitin antibodies shown to have a preference for polyubiquitin chains under denaturing PAGE-western blotting conditions revealed only modest co-localization of polyubiquitin with SGs in both HeLa and U2OS cells (**Fig. 5A,B**)^31^. However, the co-localization with SGs was far weaker than compared to the pan-ubiquitin signal (**Fig. 1E)**, suggesting that most of the ubiquitin in SGs represents either free ubiquitin or monoubiquitylated proteins. Interestingly, increasing the length of arsenite exposure in HeLa cells (2h at 250uM) resulted in co-localization of the FK1 and FK2 immunofluoresence signal with juxta-nuclear inclusion bodies reminiscent of aggresomes (**Fig. 5B**)^32^. These inclusions were rarely seen with shorter exposures to arsenite (30min at 250uM) and were not prominently stained by the pan-ubiquitin antibody (**Fig. 5B**). Treatment of HeLa cells with the Ub-E1i prior to SG induction did not alter SG formation and the pan-ubiquitin signal was unaffected. In contrast, overall staining intensity with FK1 and FK2 antibodies was significantly reduced (**Fig. 5B**). Interestingly, while co-localization of the FK1 and FK2 signal with aggresomes disappeared completely upon Ub-E1i treatment, most of the remaining signal clearly co-localized with SGs (**Fig. 5B**). These data suggest that both FK1 and FK2 can recognize free ubiquitin under immunofluorescence conditions, especially when the cellular pool of free ubiquitin is increased by the near complete ablation of polyubiquitin conjugates upon UAE inhibition. This result further supports the hypothesis that the ubiquitin pool that co-localizes with SGs consists primarily of unconjugated ubiquitin or monoubiquitylated proteins. To further substantiate these claims, we performed immunofluorescence staining with two antibodies that specifically recognize either K48- or K63-linked poly-ubiquitin chains^33^. Indeed, the signals from both K48-and K63-specific antibodies (Apu2 and Apu3, respectively) do not co-localize with arsenite-induced SGs, but instead label the same presumptive aggresome structures seen with FK1 and FK2 antibodies in a time-dependent manner (**Fig. 5C, S2**). As expected, Ub-E1i treatment ablated both K48 and K63 signals, reinforcing the linkage specificity of these antibodies and their inability to recognize free ubiquitin (**Fig. 5C, S2**).

**Figure 5.**
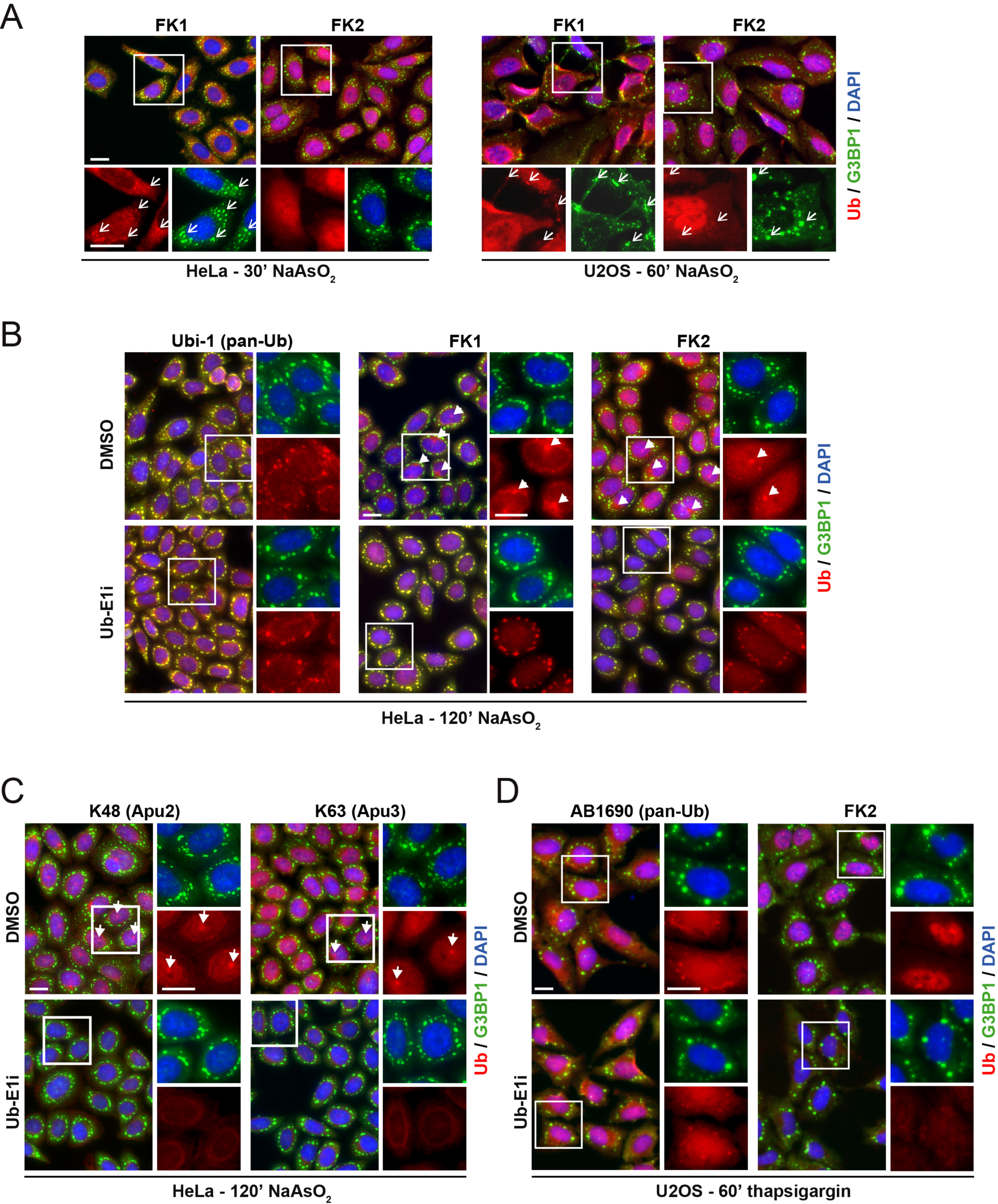
Unconjugated ubiquitin localizes to stress granules in a Ub-E1 independent manner. A).Immunofluorescence staining of HeLa and U2OS cells treated with sodium arsenite (250μM) for 30’ (HeLa) or 60’ (U2OS) prior to fixation. Cells were stained using an antibody against G3BP1 (green) and two different antibodies with a preference for polyubiquitin (FK1 and FK2, shown in red). Co-localization of ubiquitin signal with G3BP1-positive stress granules is indicated by open arrowheads. B).Immunofluorescence staining of HeLa cells pretreated with DMSO or Ub-E1i (2.5μM) for 60’, followed by treatment with sodium arsenite (250μM) for 120’ prior to fixation. Cells were stained using an antibody against G3BP1 (green) and the indicated antibodies against ubiquitin (red). G3BP1-negative juxtanuclear aggresomes are indicated by closed arrowheads. C).Immunofluorescence staining of HeLa cells pretreated with DMSO or Ub-E1i (2.5μM) for 60’, followed by treatment with sodium arsenite (250μM) for 120’ prior to fixation. Cells were stained using an antibody against G3BP1 (green) and linkage-specific antibodies against K48- and K63-linked polyubiquitin chains (red). G3BP1-negative juxtanuclear aggresomes are indicated by closed arrowheads. D).Immunofluorescence staining of U2OS cells pretreated with DMSO or Ub-E1i (2.5μM) for 60’, followed by treatment with thapsigargin (1μM) for 60’ prior to fixation. Cells were stained using an antibody against G3BP1 (green) and the indicated ubiquitin antibodies (red). Images from Ub-E1i-treated cells in panels (B), (C), and (D) were taken at the same exposure time and acquisition settings in the ubiquitin channel as images from DMSO-treated cells. Nuclei in all panels were stained using DAPI. Scale bars = 20μm in all panels.

To test the generality of our observations, we induced SG formation using thapsigargin (Tg) treatment of U2OS cells and examined pan-ubiquitin and polyubiquitin localization. Similar to arsenite-induced SGs, the pan-ubiquitin signal co-localized clearly with Tg-induced SGs and was not substantially altered by UAE inhibition in U2OS cells (**Fig, 5A,D**). By contrast, immunofluorescence staining with the FK2 antibody revealed a complete lack of co-localization between the presumptive polyubiquitin signal with Tg-induced SGs, and UAE inhibition resulted in a dramatic reduction of the FK2 signal (**Fig. 5D**). Taken together, our results clearly indicate that the ubiquitin that co-localizes with SGs is predominantly unconjugated ubiquitin and that polyubiquitylated proteins do not robustly localize to either arsenite or Tg-induced SGs. This result is consistent with our earlier demonstration that UAE inhibition does not impair SG dynamics as UAE inhibition only impacts poly and monoubiquitylation of substrate proteins. While it remains a possibility that unconjugated ubiquitin is important for SG dynamics, active protein ubiquitylation or neddylation is dispensable for SG dynamics.

## Discussion

Accumulating evidence suggests that multivalent protein-protein, protein-RNA, and RNA-RNA interactions are required to nucleate SGs and that SGs form as a result of liquid-liquid phase separation (LLPS) among SG components^34, 35^. Indeed, many identified SG proteins contain intrinsically disordered domains or domains of low complexity that are critical not only for their SG localization but also their ability to undergo phase transitions *in vitro*^36-44^. Post-translational modifications on SG proteins could serve to either disrupt critical multivalent interactions or provide new contact surfaces that drive LLPS^12^. Because ubiquitin can form polymeric chains and ubiquitin interacting proteins are present in SGs, it was conceivable that protein ubiquitylation might regulate higher order protein-protein interactions critical for SG dynamics. Combined with previous studies, our results demonstrate that ubiquitin co-localizes with SGs nucleated by a variety of protein homeostasis insults. However, our results also clearly establish that acute inhibition of active protein ubiquitylation or neddylation does not directly impact SG formation or dissolution despite the large impact on overall protein ubiquitylation. Previous studies demonstrated that long-term (18hr) NAE inhibition or knockdown of neddylation components inhibits SG formation^20, 21^. Near complete NAE inhibition using the N8-E1i takes place on minute timescales when added to mammalian cells^45^. As such, it is critical to separate observations using acute NAE or UAE inhibition with those using prolonged inhibition that will result in widespread cellular dysfunction. Our observation that ubiquitin remains co-localized with SGs even after acute UAE inhibition suggests that free, unconjugated ubiquitin may play a role in nucleating SGs. Consistent with this hypothesis is the demonstration that the ubiquitin binding protein UBQLN2 undergoes LLPS in vitro and that this transition can be inhibited by adding unconjugated ubiquitin to the reaction^46^. It will be difficult to demonstrate that free ubiquitin is critical for SG dynamics given the inability to ablate cellular ubiquitin without catastrophic consequences to overall cellular function.

Nearly all chemical SG inducers disrupt protein folding and quality control pathways resulting in the accumulation of polyubiquitylated material within cells. As such, it is likely that polyubiquitylated proteins are recruited into SGs. Evidence also suggests that defective ribosomal products, known as DRiPs, also co-localize with SGs and a relatively high percentage of DRiPs are ubiquitylated^15, 23, 27, 47^. Interestingly, heat stress-induced SGs do not immediately co-localize with ubiquitin despite early DRiP SG co-localization, as determined by OP-Puro staining^15^. Ubiquitin only co-localizes after prolonged heat stress suggesting that the recruitment of misfolded proteins results in aberrant SGs with altered physical properties^15^. The idea that SGs themselves are initially cytoprotective but only become cytotoxic upon recruitment of misfolded proteins during chronic UPS dysfunction is intriguing. The observations that the autophagy system, the proteasome, and the unfoldase p97/VCP are needed to clear these aberrant SGs are consistent with this notion^18, 19, 23, 48^. Future studies are needed to examine of the role of ubiquitin and ubiquitin pathway enzymes in regulating the dynamics and biophysical properties of these aberrant SGs.

## Materials and Methods

### Antibodies, Chemicals, and Plasmids

The following antibodies were utilized in this study. Antibodies for α-tubulin (3873), and Phospho-eIF2*α* (Ser51, 3398) were from Cell Signaling Technology. G3BP1 antibody was from MBL International (RN048PW). Antibodies against ubiquitin (AB1690, Ubi-1 (MAB1510), FK1 (04-262), FK2 (04-263)) and specific ubiquitin linkages (anti-Lys48, clone Apu2 (05-1307), anti-Lys63, clone Apu3 (05-1308)) were from EMD Millipore. Nedd8 antibody was provided by Takeda Pharmaceuticals Inc.^29^. MG132 (1748) and thapsigargin (1138) were from Tocris, MLN4924 (TAK-924, N8-E1i) was obtained from Cayman Chemical and MLN7243 (TAK-243, Ub-E1i) was obtained from Chemietek.

### Cell culture

HCT116, U2OS and HeLa cells were purchased from American Type Culture Collection (ATCC). Lenti-X 293T cells were purchased from Clontech. All cell lines were grown in complete DMEM media (Gibco) containing 10%FBS (Omega Scientific), penicillin (50 I.U./ml), and streptomycin (50 μg/ml) (Mediatech). For stable isotope labeling by amino acids in cell culture (SILAC) experiments, cells were cultured in custom DMEM without arginine or lysine (Mediatech) supplemented with 10% dialyzed FBS (Life Technologies), penicillin (50 I.U./ml) streptomycin (50 μg/ml) (Mediatech), L-Arginine hydrochloride (85µg/ml Sigma) and either “light” L-Lysine hydrochloride (50μg/ml Sigma) or heavy 13C6,15N2 L-Lysine-hydrochloride (50μg/ml Cambridge Isotopes) and 292 μg/mL L-Glutamine (Mediatech). All cell lines were grown at 37°C in the presence of 5% CO2.

### Immunofluorescence

For immunofluorescence staining, cells were fixed in 4% paraformaldehyde in PBS for 15-20’ at room temperature, rinse twice in PBS, permeabilized in 0.25% Triton-X in PBS for 15’ at RT and blocked in 5% BSA in PBS + 0.01% Tween-20 (PBST) for at least 30min at RT. Primary antibodies were diluted in blocking buffer and incubated overnight at 4°C, followed by 2 washes in PBST, counterstaining of nuclei with DAPI and two final washes in PBS. Cells were then imaged at 20X magnification on a ZEISS Axio Vert.A1 inverted microscope and images were processed using ImageJ.

### High-content imaging of SG time course plates

96- or 384-well plates were imaged using a Vala Sciences IC200-KIC high-content screening system. Either four or nine fields at the center of each well were imaged with a 20X objective through 460 nm and 535 nm emission filters for DAPI and G3BP1-GFP, respectively. Exposure times were optimized for each set of plates and applied uniformly to all wells.

### Automated image segmentation and feature quantification

Images from SG assembly and disassembly time courses were segmented and image features quantified using a custom CellProfiler pipeline^49^. Briefly, nuclei were segmented and identified in the DAPI fluorescence channel images using a diameter cutoff of 9-80 pixels for HEK293xT and HeLa cells. Cell bodies were then extrapolated by overlaying the GFP fluorescence channel images and tracing radially outward from the nuclei to the limits of the cytoplasmic G3BP1-GFP signal. The cell bodies were used as masks to eliminate imaging artifacts outside of cell boundaries, such as background fluorescence or dead cells. After masking, punctate structures were enhanced by image processing for speckle-like features that were 10 pixels in diameter for HEK293xT and HeLa cells, and these punctate structures were then annotated as stress granules. Finally, feature count and total feature area for nuclei and stress granules were calculated and exported for statistical analysis. To quantify SG dynamics, the total area covered by SGs was normalized by the total are covered by nuclei for each acquired image. Each experimental condition was represented by between 12 and 16 replicate wells, with between 4 and 9 images acquired per well. The mean and standard deviation were calculated across all replicate wells and fields of view acquired per well, with a minimum of ∼25,000 cells analyzed per condition.

### Mass Spectrometry

Heavy and light labeled cells were mixed 1:1 and processed for proteomics and diGLY-immuno-affinity enrichment as described previously^26, 27^. Samples were analyzed by LC-MS/MS using a Q-Exactive mass spectrometer (Thermo Scientific, San Jose, CA) and all data was processed as described previously^26^. All RAW mass spectrometry data files have been deposited at the MassIVE repository using the accession identifier MSV000082933.

## Author contributions

Conceptualization, S.M. and E.J.B.; Methodology, S.M., G.W.Y. and E.J.B.; Formal Analysis, S.M., A.F. and E.J.B.; Investigation, S.M., A.F. and R.H.; Writing – Original Draft, S.M. and E.J.B.; Writing – Review & Editing, S.M., E.J.B.; Visualization, S.M. and E.J.B.; Supervision, G.W.Y. and E.J.B.; Funding Acquisition, G.W.Y. and E.J.B.

## Declaration of Interests

G.W.Y. is a co-founder of Locana and Eclipse Bioinnovations and member of the scientific advisory boards of Locana, Eclipse Bioinnovations and Aquinnah Pharmaceuticals. The terms of this arrangement have been reviewed and approved by the University of California, San Diego in accordance with its conflict of interest policies. All other authors declare no competing interests.

## Supplementary Information

**Figure S1.**
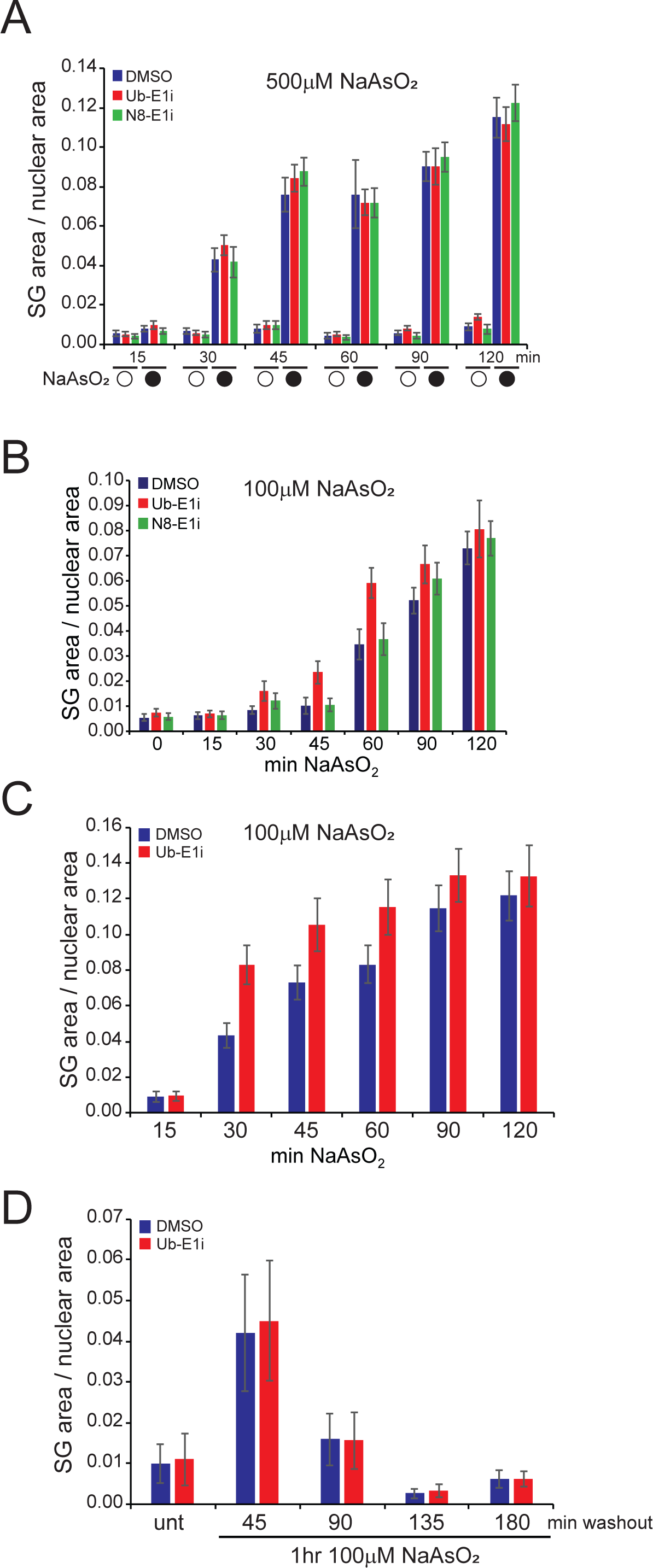
Stress granule dynamics are unaffected by inhibition of protein ubiquitylation or neddylation. A).293T cells expressing GFP-tagged G3BP1 were pre-treated for 90’ with DMSO (blue bars), Ub-E1i (red bars), or N8-E1i (green bars) followed by treatment with (black circles) or without (white circles) sodium arsenite (500μM) for the indicated times. Cells were fixed and imaged at the indicated times. B).293T cells expressing GFP-tagged G3BP1 were pre-treated for 90’ with DMSO (blue bars), Ub-E1i (red bars), or N8-E1i (green bars) followed by treatment with sodium arsenite (100μM) for the indicated times. Cells were fixed and imaged at the indicated times. C).Hela cells were pre-treated with DMSO or Ub-E1i for 90’ followed by treatment with sodium arsenite (100μM) for the indicated times. Cells were fixed and SGs were localized using G3BP1 immunofluorescence. D).Hela cells were pre-treated with DMSO or Ub-E1i for 90’ followed by treatment with sodium arsenite (100μM) for 60’ at which time the sodium arsenite was washed out and cells were fixed at the indicated times following sodium arsenite washout. SGs were localized using G3BP1 immunofluorescence. For all panels, the stress granule and nuclear area was quantified using custom image analysis scripts for the indicated time points and treatments. Error bars depict standard deviation of the mean.

**Figure S2.**
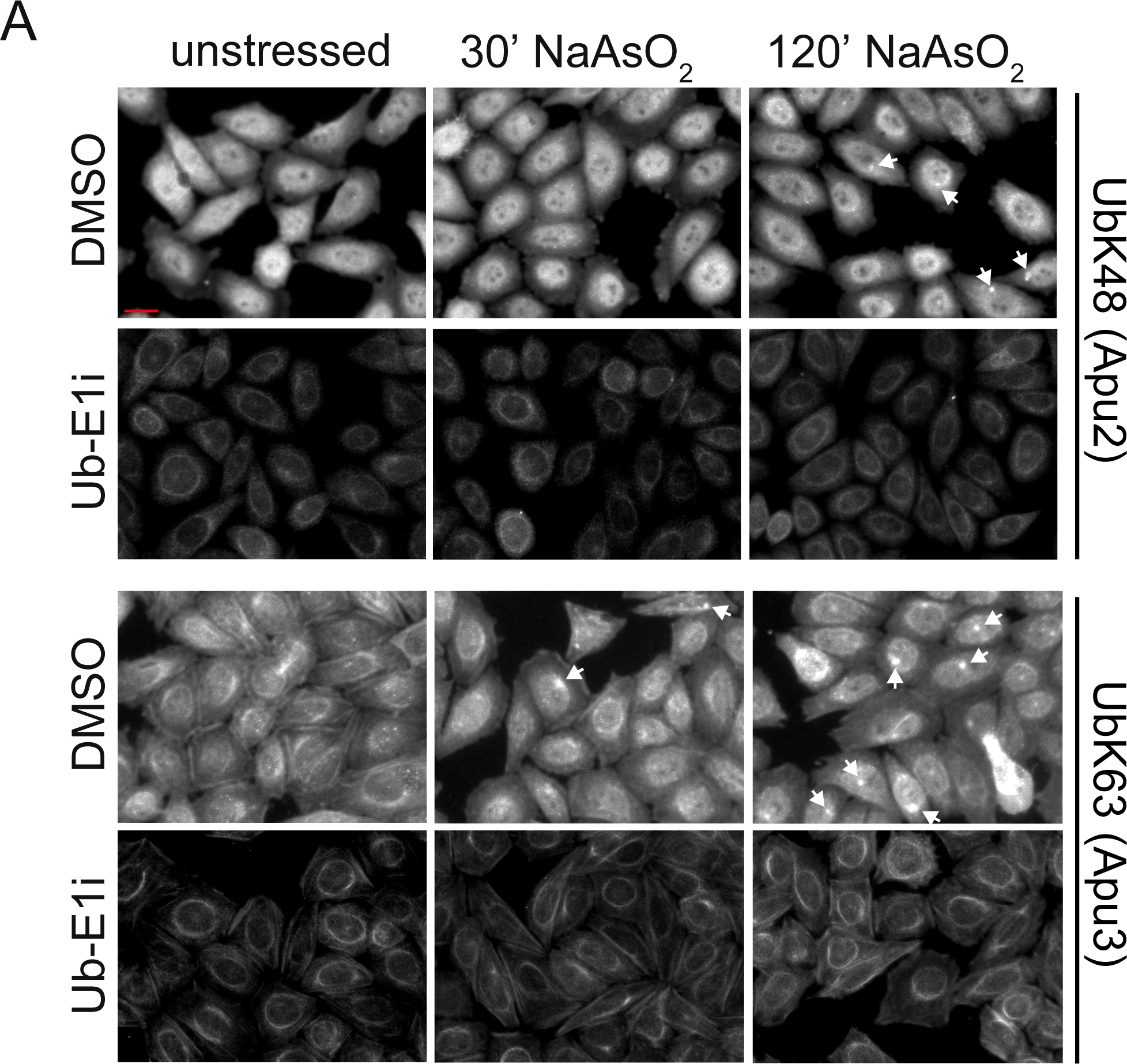
Lysine 48 and 63-linked polyubiquitin chains do not co-localize with SGs. Immunofluorescence staining of HeLa cells pretreated with DMSO or Ub-E1i (2.5μM) for 60’, followed by treatment with or without sodium arsenite (250μM) for the indicated times prior to fixation. Cells were stained using linkage-specific antibodies against K48- and K63-linked polyubiquitin chains. Juxtanuclear aggresomes are indicated by closed arrowheads. Scale bar = 20μm.

**Table S1. – Alterations to the proteome and ubiquitin-modified proteome upon arsenite treatment and washout.** Table of identified proteins and diGLY-modified peptides and associated SILAC (H:L) ratios from HCT116 cells treated with sodium arsenite (500μM) for 0’, 20’, 45’, or treated for 45’ followed by a 3hr washout. Heavy labeled cells were arsenite treated.

**Table S2. – List of annotated stress granule-localized proteins.** The curated list of previously determined SG-localized proteins used for all proteomic analysis of known SG proteins.

**Table S3. – Alterations to the proteome and ubiquitin-modified proteome upon UAE inhibition.** Table of identified proteins and diGLY-modified peptides and associated SILAC (H:L) ratios from HCT116 cells treated with either the Ub-E1i (1μM) alone or the Ub-E1i in combination with MG132 (10μM) for 4 hours. Heavy labeled cells treated with the Ub-E1i were mixed with untreated light-labeled cells and heavy-labeled cells treated with the Ub-E1i and MG132 were mixed with light-labeled cells treated with MG132 alone.

